# Relating farmers’ perceptions to long term dynamics in maize (*Zea mays* L.) production: The production of improved maize is coming to a standstill in central Malawi

**DOI:** 10.1101/2020.11.02.364604

**Authors:** Harrington Nyirenda, Wantwa Mwangomba, Ellen M. Nyirenda

## Abstract

Maize production, area and yield dynamics were assessed based on farmer perceptions and production data from 2004/05-2018/19 using 36 000 households in Salima, central Malawi. The results showed that farmers used 16, six and two varieties for hybrid, Open-pollinated varieties (OPV) and local maize respectively. Farmers sourced Hybrid and OPV maize seed from Private Agro dealers while local maize was own-sourced. Farmers preferred local maize for being cheap, good taste, low storage costs, and pest resistance although low yielding. They preferred hybrid and OPV maize for high yielding and early maturity despite demanding high storage costs, pest susceptibility, and low flour. From 2004/05-2018/19, the area under local and OPV maize reduced by 61% and 12% respectively, while that of hybrid maize increased by 49%. However, the consistent decrease in area for hybrid and OPV and significant increase of that of local maize from 2014/15-2018/19 may signal a catastrophic maize production in the region. From 2019/20-2025/26 production of all maize was projected at 44 172 tons by 2025/26, representing a 1.6% increase from the base year 2019/20. This increase will be due to favorable climatic conditions and not increase in area or yield. If maize yield was improved by 30% production would increase to 110 430 tons representing 67% of the food requirement in the study area. The current maize production trend in Salima does not guarantee food security prospects. Therefore, policymakers should consider reviewing the past interventions (input pricing, promotion strategies, sustainable practices, policies) in the maize subsector to enhance maize productivity.

## Introduction

Food self-sufficiency in Malawi largely depends on the amount of maize harvested in a season (FAO 2015). The dominant subsistence agriculture is characterized by low yield, with a national average of about 1.6 t ha^−1^ (FAO 2020) despite the promotion of the use of improved maize varieties. Maize production is predominantly rain-fed, practiced once (November-April) in a season, although some farmers practice irrigation.

Poor crop performance has been attributed to dependence on rain-fed production, limited use of improved seeds and fertilizers, climate change, degraded soils, limited markets, and inadequate agricultural extension system (Government of Malawi 2011; FAO 2015). In 1992, the average national soil loss was estimated at 20 t ha^−1^ y^−1^ (World Bank 1992) while in 2016, the figure had risen to 29 t ha^−1^ y^−1^ with Salima District registering 7.6 t ha^−1^ y^−1^ (Vargas and Omuto 2016). This soil loss is estimated to have led to the loss of topsoil worth valuable nutrients useful for maize production. In some studies, the annual loss had been estimated at USD2 billion soil nutrient equivalent (Chigowo 2007). The country experiences high effects of climate change and variability especially along the hot areas of Lakeshore and extreme south. Droughts and floods have alternated in successive seasons. These have wiped out near-maturing maize or dried up the crop in the fields. With limited irrigation capacity, and failure to meet required production during the rainy period, hunger has been felt in most communities. Low yields mean that farmers do not adequately sell their crop which limits their financial capacity to purchase adequate inputs for the subsequent seasons. In good seasons, when farmers realize bumper harvests, farm gate prices are usually low (Chirwa 2006; Tchale and Sauer 2007). The ‘time to wait’ for higher prices is affected by the need to meet crop storage costs (Edelman et al. 2015). With limited financial means, the post-harvest loss is high among most subsistence farmers. Food shortage at the household level results in some socioeconomic crises such as theft and unprotected sexual activities (leading to contraction of HIV/AIDS, unplanned pregnancies, abortion) among girls and women (Conroy 2006). These issues led to the advancement of a policy direction aiming at improving maize yield and productivity (Levy 2005). For example, in the 2005/06 season, the Government of Malawi introduced the Farm Input Subsidy Programme (FISP). The program aimed at increasing maize yield and production on a small piece of land using improved maize varieties and the application of enough fertilizers (Government of Malawi 2018a). The policy was a direct response to the famine of the 2004/05 season where 4.2 million Malawians needed food aid (Malawi Vulnerability Assessment Committee 2005.). FISP has run for 13 years continuously (still on). The design is that a household is provided with improved maize seed and fertilizers for a 0.4 ha. The program has gone through several adjustments along the period such that at the national level, 1.5 million households benefited in 2005/06 and the number reduced to 900,000 households in the 2018/19 season. In 2005/6, the country achieved a surplus of about 510 000 tons (above national requirement) with a yield of approximately 1.59 t ha^−1^ while in 2018/19, the yield was at 1.62 t ha^−1^(FAO 2020). During the period of FISP, the emphasis has been placed on the use of improved varieties and the current extension message to farmers is that they should stop growing local maize varieties because of low yields and production. This is an attempt to thrive the Malawi economy which relies on agriculture production; a sector accounting for 80% of the total labour force. Its population, currently at 17.5 million with 3.5 million farming households (Government of Malawi 2018b), needs sustainable production mechanisms (Government of Malawi 2006).

The present study intended to fill two research knowledge gaps. Firstly, information is scanty on the status of local, hybrid, and Open-pollinated variety (OPV) maize with respect to yield, production, and area over a long period (about 15 years) in most agricultural divisions in Malawi. OPV maize seed can be recycled to a maximum of three years while hybrid seed can not be recycled (Denning et al. 2009; Kutka 2011). There are claims that improved maize varieties provide sustainable production trends and meet farmers’ needs (Government of Malawi 2006) but systematic assessment to ascertain these claims under farmer field conditions and farmer perceptions for a longer period has not been done in respective agricultural divisions in Malawi. It has been claimed that most of the hybrid varieties do not meet Malawian farmers’ desired characteristics like storability and poundability (Lunduka et al. 2013; Fisher et al. 2015). Such studies have not reflected the trend in local and hybrid maize production over a longer period to compare farmer perceptions and field practice trends. Questions could still be asked as to whether hybrid or improved maize alone would meet food requirements and other perceptions of farmers for the ever-increasing population trend in maize growing regions like Malawi. Most claims are based on on-station and on-farm research conditions (Magorokosho et al. 2010; Setimela et al. 2012) which lack adequate replications of farmer conditions. Other studies are based on a very short period e.g. one to five seasons like those of Smale et al. (1991) for the 1989-1990 season, Denning et al (2009) for 2005/06-2006/07, and Holden and Fisher (2015) for 2006/2009-2011/2012 seasons only. Some research has also been based on low sample sizes, for example, in Setimela et al. (2012), only 49 households in eight (8) African countries were sampled with a focus on lead farmers only. The study by Holden and Fisher (2015) covered the 2011/12 season and sampled 350 farming households from six Districts in Malawi (possibly 58 households per district). Much as studies of that nature may capture wider scope on agro-ecologies, they may present deficiencies on long term production dynamics which may hinder prediction prospects.

Secondly, maize production estimates in Malawi do not take into consideration the predictive factors, rather, it has been based on the past production trend (Ministry of Agriculture and Food Security 2008). With the increase in population in Salima Agricultural Development Division (SLADD) (Government of Malawi 2018b), the consumption of maize food products will obviously increase. As such, the estimation of supply and production, in particular, is extremely important in the context of long and short term planning to achieve and improve food security development objectives (Government of Malawi 2006). Projecting the production of the maize crop is important to identify possible gaps in supply, especially in Malawi where maize is the main staple food, to avert the risks of food insecurity. Such projection estimates may improve decision making regarding input-factors that affect the input supply sector and agribusiness on the part of the government and may influence farmers’ decisions on the allocation of input to particular food crops (Meyer 2005). A comprehensive projection of maize production levels has not been undertaken for SLADD, therefore, there is a need to project the future production level of maize commodity for at least the next five year planning period that will feed into future demand and supply balance policy interventions. This will also feed into the District Development Plans (DDPs) formulated every five years.

To understand the two problem areas above, the present study focussed on one of the agricultural divisions; Salima Agricultural Development Division (SLADD), particularly in Salima District. Our long term study provided a suitable regime to understand farmers’ perceptions, area cultivated, yield, and production trend. The duration allowed further understanding of percentage changes of the parameters and provided a basis for prediction. The long term study also allowed for a better understanding of maize production dynamics even during good or calamitous seasons like pest outbreaks such as African armyworm and Fall armyworm. The Fall armyworm was first observed in Malawi in 2016 (Day et al. 2017).

The study used both qualitative methods i.e. focus group discussions (FGD) with community members, direct field observations (DFO), and quantitative i.e. time series data on maize production, yield, area, local maize price, local rice price, fuel data (a proxy for input costs), rainfall levels.. The combined participatory methods reduce limitations met in conducting traditional research methods (relying on yield, production data only) (Kumar 1987. These methods enable joint understanding between the etic (scientist as outsider-in this case) and emic (farmer as insider-in this case) (Prince 2001; Guimaraes and Mourao 2006). Kumar (1987) defined group interviews as the use of different direct techniques in probing to collect information from people in a group set up of varying compositions or sizes. The probing can be done by one or more people through the use of a guide or not. In this paper, hybrid and OPV are also referred to as improved maize.

The study attempted to answer the following research questions:

1. What are the common varieties of hybrid, local, and OPV maize grown in the Salima Agricultural Development Division?
2. What factors do farmers consider when making a decision to choose between a local and improved maize?
3. What is the extent of cultivation of hybrid, local, and OPV maize in terms of area, yield, and production among farmer conditions in Salima Agricultural Development Division?
4. What is the maize production outlook seven years after the year 2018/19 in Salima Agricultural Development Division

## Materials and Methods

### Ethics statement

#### a) Ethics committee clearance

The researchers did not get a special clearance from the Ethics Committee because the institution where the researchers work does not have a policy requirement limiting research studies due to non-approval. Moreover, the insitution does not have an Ethics Committee for research studies. Researchers just inform the controlling officers in respective duty stations about their day today scheduled activities (routine activities) and authorities approve those routine activities. This was the case with us. This study was part of routine work.

#### b) Approval from participants

Researchers got an approval from the chiefs for the involvement of farmers in focus group discussions. This is according to the tradition. The approval was requested and obtained orally. The chiefs and researchers obtained oral approval of participation in the focus group discussion by farmers.

### Study area

The study was done in Salima Agricultural Development Division (SLADD), particularly in Salima District. SLADD comprises two districts of Salima and Nkhotakota. Salima district is located at Latitude 13° 47 S, Longitude 34° 26 E, central Malawi. The distinct seasons include warm wet (November to April), cool dry (May to August), and hot dry (September to October) (Unpublished data, Malawi Meteorological Office). The district has a total area of 597 588 ha (107 377 ha is cultivatable; 12 351 ha under irrigation; 6 888 ha of forest reserves; 56 146 ha under the estate sector). The district population was estimated at 478 346 with a projection of 548 346 by 2026 (Government of Malawi 2018b). Landholding size among smallholder farm households ranges from 0.4 to 0.8 ha while estates range from 10 to 400 ha. The district had 123 396 farming households. An average farm household had about 4.4 members with an annual per capita consumption of 300 kg of maize. Rainfall (Figure 1) is unimodal (October to April) and ranges from 725 mm to 2 500 mm. This range is characterized by either too much in a short time or little in an extended period (erratic). Temperature ranges from 20°C (minimum) to 30.7 °C (maximum) with October as the warmest and July as the coolest months respectively. The district is dominated by Chromic, Haplic, and Luvisols soil types, with a depth of up to 150 cm (Lowole 1995). However, Calcimorphic alluvial (Inceptisol) and Vertisol, dark fertile soils, are also found in some parts of the district (Saka et al. 2003). The dominant crops under smallholder farmers include maize (*Zea mays*), groundnut (*Arachis hypogaea*), rice (*Oryza linu*), sweet potato (*Ipomoea batatas*), and cassava (*Manihot esculenta*) whereas cash crops include tobacco (*Nicotiana tabacum*), beans *Phaseolus vulgaris*, cotton (*Gossypium hirsutum*), soya (*Glycine max*), sorghum (*Sorghum bicolor*), cowpea (*Vigna unguiculata*) and chilies (*Capsicum frutescens* L and *Capsicum annuum* L.) (Beedy et al. 2015; Salima Agricultural Development Division 2019).

**Figure 1.**
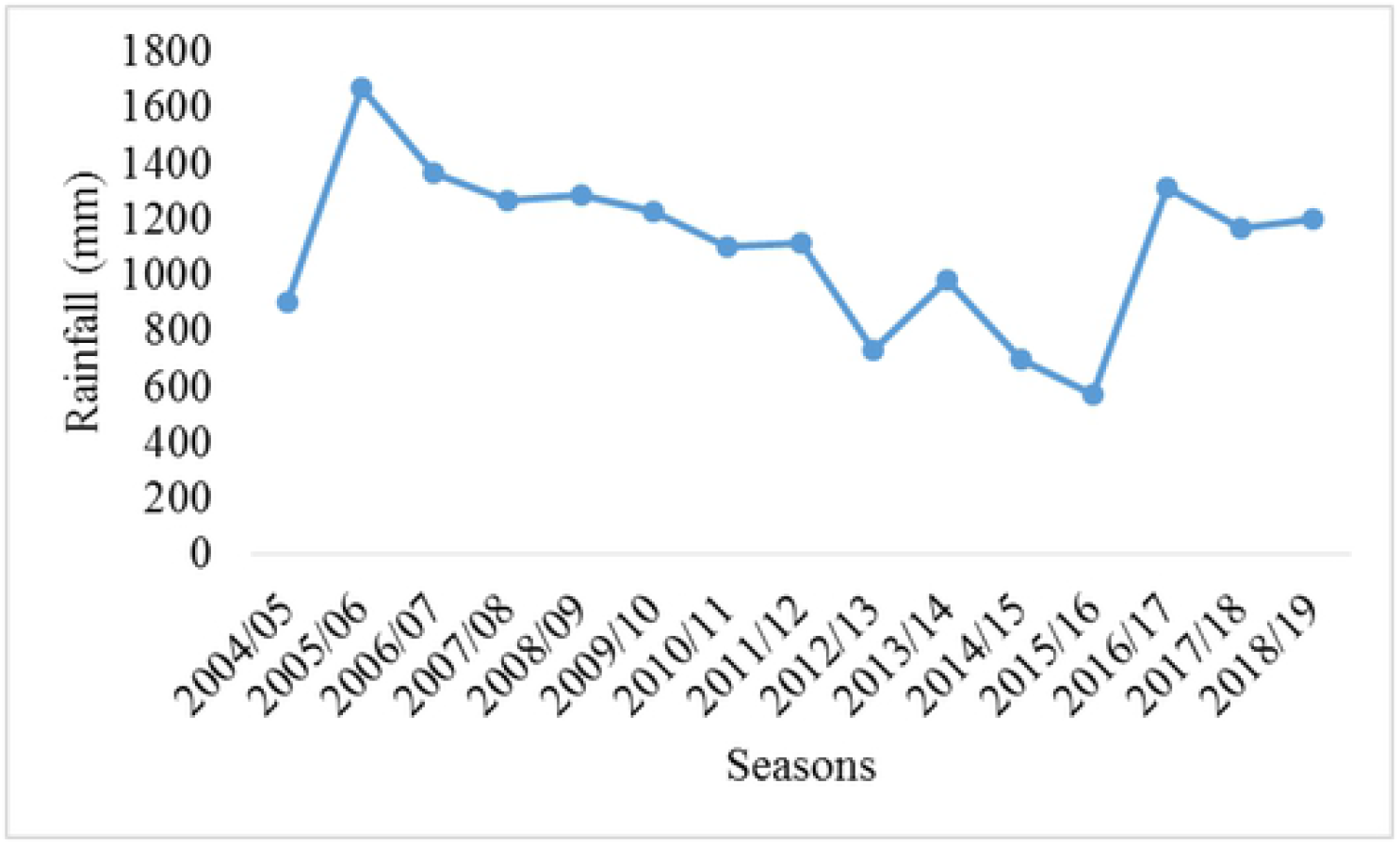
Rainfall in the study area for the period of the study. Source: Department of Climate Change and Meteorological Services, unpublished data

### Data collection

#### Farmers’ perception of local, hybrid, and OPV maize production (Focus group discussion)

To understand farmers’ perception of the use of local and improved maize varieties, this study followed Constructivism Epistemology (CE) approach (Sayre 2001). This entails the development of knowledge-based on active interaction and involvement of research participants e.g. community members and the researchers. Participants provide information based on a defined topic they have knowledge and expertise on (Guerci and Shani 2014). CE methods include case studies, focus group discussions (FGDs), participant observation, and phenomenological study. In our case, we chose FGDs compared to general community interviews because the latter are dominated by few leaders hence locking up some information from others; chorused responses can also interfere with the real information being sought (Kumar 1987; Elias 2013). During FGD, with the help of a moderator, members are free to express views on their ideas, issues, experiences, and insights among one another (Kumar 1987). Any member can elaborate, criticize, and comment on views expressed by another. This provides an opportunity to elicit elaborate information. An appropriate FGD is guided by a research question to ensure study replicability, reliability, and validity (Chioncel et al. 2003). A research question informs and guides the FGD. In our case, the research question was ‘ *What factors do farmers consider when deciding on the type of maize to grow?* To ensure stability and focus on the topic, the major discussion points were formulated together with participants based on their experience and knowledge (Dansoh et al. 2020). A total of 15 individual farmers were purposefully selected to represent major categories of farmers on maize production. A valid FGD uses participants with competent knowledge on the research topic (Chioncel et al. 2003) therefore, the participants had members characterized by (i) having done maize production (local or/and improved maize) for not less than 10 years (ii) Lead farmers in maize production. The FGD was held in local settings to ensure free, relaxed-interaction between researchers and participants (Dansoh et al. 2020). The FGD lasted for 3 hours (Kumar 1987) with two 20 minute breaks in between. The FGD was guided by an experienced moderator, with knowledge of the study topic, group dynamics, and local languages. The rest of the researchers were taking notes and observing participants’ reactions.

Direct Field Observations (DFOs) were also done jointly with members involved in FGD to appreciate the farmers’ fields with either local or improved varieties. A total of 10 maize fields were visited and observations were made. DFO complemented the group interviews. This helped get the actual practices. The DFOs enhanced understanding of the descriptions made during the interviews; all or some omitted/forgotten issues during the interviews could be captured during the DFO. DOFs widen understanding as some group responses are sometimes just ‘chorused’ without substantiations (Orr and Richie 2004; Sileshi et al. 2007).

#### The extent of local, hybrid, and OPV maize production (Agriculture production estimate data)

To understand the production dynamics, data were collected on area cultivated, yield, and production for 15 seasons (2004/05-2018/19) for local, hybrid, and OPV maize following the Agricultural Production estimate guidelines (Ministry of Agriculture and Food Security 2008). The district was divided into 80 sections, each divided into eight blocks. Each season two (2) blocks per section were selected using a random sampling number. Then all the farming households from the two randomly selected blocks were listed separately. In each of the two selected blocks, 15 farming households were sampled using the random household number. Therefore, a total of 2 400 households were sampled each season to estimate maize production for the district (36 000 households in 15 years). To ensure we captured the existing dynamics, data collection was done on the default sizes of maize fields/plots (Holden and Lunduka 2012) of either local or improved maize of the sampled households. The default maize field/plot where measurements were done was called Yield Sample Plot (YSP). Various numbers of YSPs were studied in a season depending on the number of maize field/plots a sampled household had. Major parameters of varieties used, area, yield, and production were recorded. The maize field/plot area was measured using a GPS. Disastrous events like floods, droughts, and pest attacks (Fall armyworm, African armyworm) were also noted. Each season new households were sampled to ensure representativeness of spatial coverage and production dynamics.

For maize production outlook from 2019/20-2025/26 seasons, the primary data above was supported by secondary data obtained from various sources such as Salima Agricultural Development Divisions (SLADD), Agricultural Development and Marketing Cooperation (ADMARC), Department of Climate Change and Meteorological Services (DCCMS), the National Statistics Office (NSO) and other related agencies in Malawi.

### Data analysis

#### Perception of farmers on local, hybrid, and OPV maize (FGD)

The data were analyzed using Epistemological Orientation (EO) (Sayre 2001) and Content Analysis (CA) (Krippendorf 2004) on social constructivism. EO and CA have a long history of analysing FGD data since the 1960s (Harris 2001). The views in these theories tend to emphasize how group members collaborate on some issues, how they achieve consensus (or fail to), and how they construct shared meanings about commercial products, communications, or social concerns. Specifically, the analysis followed the Pragmatical CA and Semantical CA, The meaning was derived as follows: (a) the frequency with which an idea appeared, was interpreted as a measure of importance, attention, or emphasis; (b) the relative balance of favourable and unfavourable attributions regarding a symbol or idea, was interpreted as a measure of direction or bias; and (c) the kinds of qualifications and associations made with respect to a symbol or idea, was interpreted as a measure of the intensity of belief or conviction (Krippendorf 2004). In some cases, direct quotes of farmers were noted and maintained for reporting purposes. The issues raised by the participants were summarised and laid as a transcript (Krueger and Cassey 2009). A draft transcript supported by notes was read by all researchers to confirm consistency. Then it was shared with five participating farmers as a pre-test of a common understanding of the issues (Morrisey and Higgs 2006). The farmers provided their comments and a final report of results was prepared.

#### Maize production estimate data

Data for the area, yield, and production were tested for normality (Shapiro-Wilk normality test) and homoscedasticity (Levene test). Yield data met normality and homoscedasticity while area and production data met homoscedasticity. One away analysis of variances (ANOVA) was performed to understand differences in area cultivated, yield, and production among hybrid, local, and OPV maize. A regression analysis was performed to determine the effects of rainfall on the yield of the three maize categories. A projection was performed to estimate the extent of food maize production against the food requirement in the study area. This projection was based on the mean area, production, and yield of all maize combined because the aim was to determine the food security situation. Since maize production estimate requires that a recent five year period be presented and analysed for informed comparison (Ministry of Agriculture and Food Security, 2008), we also did a thorough analysis of the 2014/15-2018/19 seasons and a projection from 2019/20-2025/26. R Statistical Package Version 3.6.2 (R Development Core Team 2020) was used to compute the data.

#### Maize production outlook from 2019/20-2025/26 (projection)

The time series covering the period from 2004/05 to 2018/19 were employed in this study. The area, production, and yield for local, hybrid, and OPV were respectively pooled for the purposes of prediction since all the maize categories are considered as one package when estimating food security (Ministry of Agriculture and Food Security 2008). The Nerlovian Partial adjustment model (Haile et al. 2016) was applied to estimate the response and project the production level of maize commodity in Salima. This supply model implies that changes in area harvested are in the proportion to the difference between the long-run equilibrium area and an actual area in the previous year. Under this methodological approach, acreage (area harvested) – one of the major drivers of maize grain production - is expressed as a partial adjustment function with the current price of maize and the prices of rice commodity which is the second cereal food to maize in Salima. It is also assumed that farmers’ decisions are based on the food prices of the previous growing season (Mapila et al. 2013). Projecting the decision on acreage can indicate how much maize will be available in the subsequent harvesting seasons. As Haile et al. (2016) put it that the amount of land allocated to maize production acts as a sound indication of food availability which has significant implications for food security situations in the country. In other words, with the yield of the maize crop available projecting the area under crop cultivation provides an important step into projecting the production of maize crop. In addition, the derived elasticities of maize crop acreage will provide the extent to which maize production responds to changes in endogenous factors.

Various equations were estimated to represent the supply model of the maize commodity in Salima. The equations were estimated using historical time series data from 2004 to 2018 of both endogenous and exogenous model variables. The outcome and reliability of the model were based on the *priori* and sound coefficients. The assumptions of Constant Returns to Scale (CRS) and perfect competition were relaxed a bit considering the size of the sample. The total supply-side equation of the maize market consists of the area, yield, production, and opening stock equations (Parappurathu et al. 2014). Various factors such as rainfall, costs of inputs, technological progress (improved varieties of maize seeds, irrigation, fertilizer application, and others that affect farmers’ decision making during the production of maize) are also considered in the maize commodity supply model. In this supply model, the production equation is considered as an identity which simply applies its formula as:

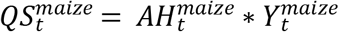

Where,

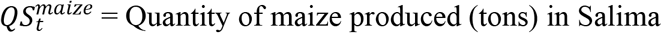
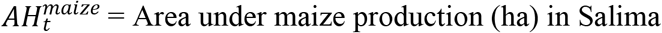
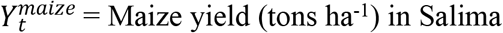

We modeled the area under production as a function of the domestic (ADMARC) price of maize for Salima district, the lagged area, and input costs (the price of fertilizer) using the Ordinary Least Squares (OLS) estimation method. For purposes of robustness, several functional forms were estimated as reported in Table 1 below however, model (2) was considered for forecasting the production of maize owing to its conformity of economic theory and statistical properties. In economic theory, the previous area under maize production, the current price of maize commodity, and the input costs affect the decisions the farmers make towards production (Haile et al. 2016). The expectation was that the lagged area and domestic maize grain price had positive relationships with the area considered for production in the current growing season, while the local price of rice – the second cereal commodity – had a negative relationship. This, therefore, gave the area equation as expressed below:

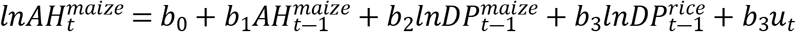

**Table 1.**
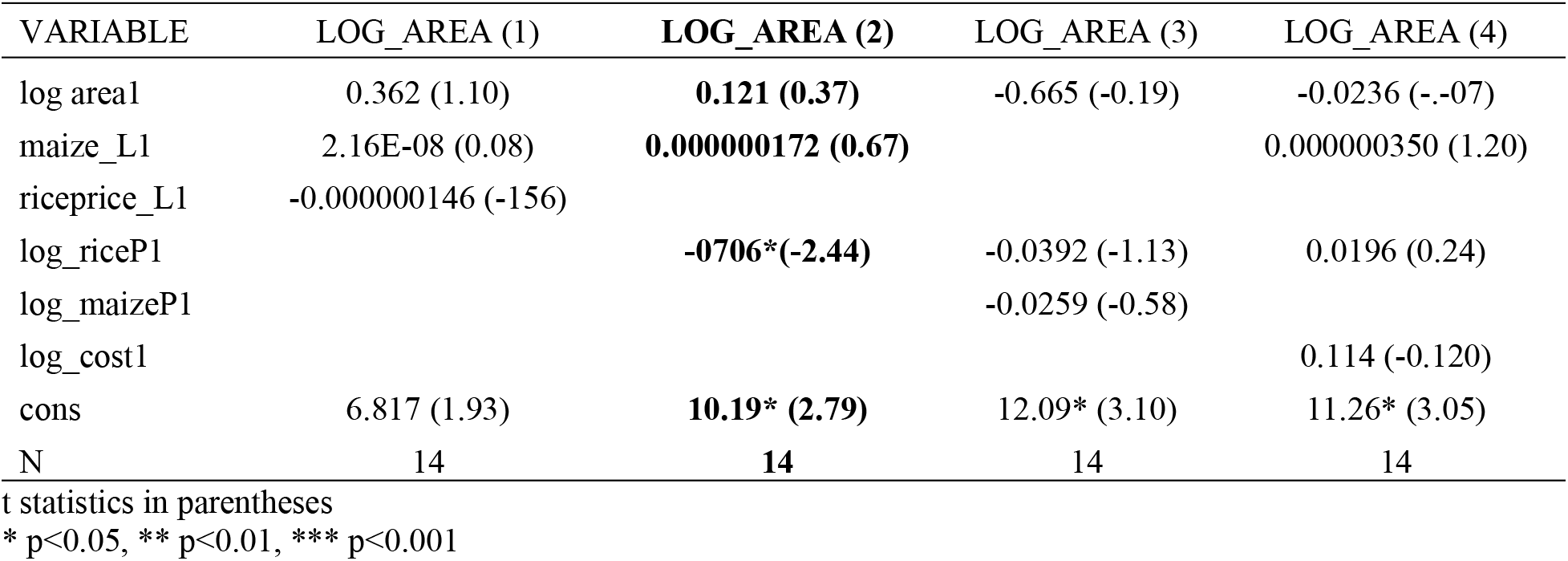
Area under maize production Equation Estimates in Salima Agricultural Development Division, Salima District, Central Malawi

| VARIABLE | LOG_AREA (1) | LOG_AREA (2) | LOG_AREA (3) | LOG_AREA (4) |
| --- | --- | --- | --- | --- |
| log area1 | 0.362 (1.10) | **0.121 (0.37)** | -0.665 (-0.19) | -0.0236 (-.07) |
| maize_L1 | 2.16E-08 (0.08) | **0.000000172 (0.67)** |  | 0.000000350 (1.20) |
| riceprice_L1 | -0.000000146 (-156) |  |  |  |
| log_riceP1 |  | **-0706\*(-2.44)** | -0.0392 (-1.13) | 0.0196 (0.24) |
| log_maizeP1 |  |  | -0.0259 (-0.58) |  |
| log_cost1 |  |  |  | 0.114 (-0.120) |
| cons | 6.817 (1.93) | **10.19\* (2.79)** | 12.09* (3.10) | 11.26* (3.05) |
| N | 14 | **14** | 14 | 14 |

t statistics in parentheses
\* p<0.05, \*\* p<0.01, \*\*\* p<0.001

Where,

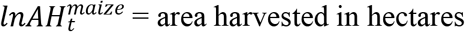
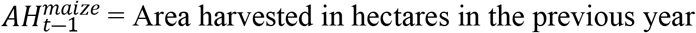
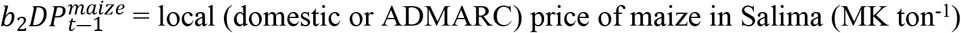
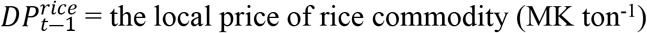
i.e. MK = Malawi Kwacha (Currency for Malawi)

Whilst the yield function of the maize supply model is estimated based on rainfall and its squared term, the natural log of the trend which captures the technological advancement and also on per capital is also considered (Meyer 2005). In Malawi, maize yield is predominantly determined by good previous year rainfall (t-1) (Mapila et al. 2013) hence a presumptively positive relationship with the yield. The maize yield equation also estimated that beyond some optimal level, rainfall begins to influence negative marginal productivity yields. The yield equation adopts model 1 in Table 2 which conforms to the economic priori and statistical properties of significance.

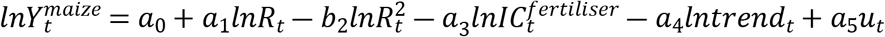

**Table 2.** Maize yield Equation Estimates in Salima Agricultural Development Division, Salima District, Central Malawi

| VARIABLE | LOG_YIELD (1) | LOG_YIELD (2) | LOG_YIELD (3) |
| --- | --- | --- | --- |
| log_R | 0 (.) | 0 (.) |  |
| log_aqr_R | 0.0344 (0.36) | 0.0312 (0.27) | 0.0312 (0.27) |
| log_cost | -0.370* (-2.24) |  |  |
| log_trend | 0.444* (2.66) | -0.351 (1.25) | 0.351 (1.25) |
| log_cost1 |  | -0.302 (-1.48) | -0.302 (-1.48) |
| _cons | 1.459 (0.88) | 1.241 (0.75) | 1.241 (0.75) |
| N | 15 | 14 | 14 |

t statistics in parentheses
\* p<0.05, \*\* p<0.01, \*\*\* p<0.001

Where,

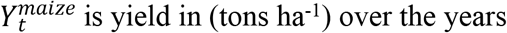
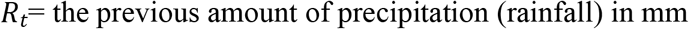
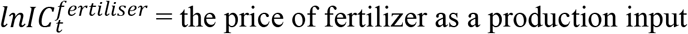
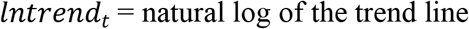

The production of maize in this paper is projected based on area response and yield functions.

## Results

### Farmers’ perception of local and improved maize

The focus group discussion resulted in the generation of six main discussion components or ‘knowledge areas’ regarding farmers’ perception of improved and local maize. The components included: *Local and improved maize varieties used by farmers*; *Source of maize seed*; *Attributes of a good maize variety or type*; *Weaknesses and strengths of local and improved maize*; *Special focus on poundability*; and *Promotion priority between local and improved maize*. The components and perceptions of farmers are outlined below:

### Local and improved maize varieties used by farmers

Participants outlined a number of maize varieties that were grown in the study area. The varieties under hybrid maize included: SC 627 (*Mkango*), SC 403 (*Kanyani*), SC 719 (*Njovu*), SC 407, DK 9089, DK 8053, DK 8033, DK 777, MH 18, MH 19, MH 26, MH 31, MH 41, PAN 4M-19, PAN 53, Pannar 413. The farmers mentioned the following as OPV varieties: ZM 421, ZM 521, ZM 523, ZM 621, ZM 623, Chitedze 4. The local varieties included *Kaduya* and *Bantam*.

### Source of maize seed

Under this component, participants identified six major sources of maize seed. For improved seed, farmers indicated (in order of dominance) that they bought from stable Private Agro dealers, ADMARC (a government institution), and mobile vendors. Some farmers received from their relatives who worked in towns and cities while others got the free seed from Non-Governmental Organisations. Depending on the variety of improved seed (hybrid or OPV), some could get free subsidy seed from Agro dealers and ADMARC or pay a small top-up amount of money (this applied to those who benefitted from FISP). Local maize seed was sourced locally. They expressed that there were no markets for local maize seed in the study area. On this point, participants indicated that they either kept their own local maize seed through selection for next season or requested from friends, relatives, or neighbors. They indicated that it was usually provided free of charge by friends or relatives. However, it was indicated that some farmers recycled hybrid seed because they could not afford the annual purchase of new seed. The debate elicited thought-provoking points for seed researchers, policymakers, or maize scientists especially on the need to ensure farmers’ participation in seed growing and managing the whole process themselves. They indicated that they would prefer to handle the seed process by themselves rather than buying every year. An interesting point was uttered as follows:

> ‘We are discouraged from processing hybrid seed (plant, select, and store) at farmer level for next seasons. We are advised to buy new seeds every season. Where is the element of sustainability with this hybrid maize? I feel buying every season is like feeding hand to mouth and we are making certain groups of people or companies rich every year. I like most of the good attributes of hybrid maize but I do not like the condition of buying seed every season’. [Lead farmer]

### Attributes of a good maize type or variety

Farmers were asked to list what they felt were a positive attribute of a maize type or variety. On this component, a number of attributes were outlined by the participants. In summary, farmers would like to have maize variety that has *good poundability*, *high yielding, cheap, readily available, good taste, adequate flour after milling, early maturing, demanding less or no fertilizer use, pest, and disease resistance in field and storage*. When asked to rank these attributes, the participants took some extended time to do the preference. This could mean diversity and differences in personal needs. Therefore, some traces of personal compromises could not be ruled out. The priority list was as follows: 1) High yielding, 2) adequate flour after milling, 3) cheap, 4) pest and disease resistance in field and storage, 5) good taste, 6) good poundability, 7) early maturing, 8) demanding less or no fertilizer, and 9) readily available.

### The weaknesses and strengths of local and improved maize

The aim was to capture perceptions as the basis for the proliferation of either local or improved maize. It must be indicated that in some cases it was not straight forward in differentiating hybrid from OPV maize by some farmers. However, some clear differences were noted when examples were outlined for specific maize variety attributes. This made the analysis of results clear. The most important aspect was that all farmers involved had a clear knowledge of the differences between local (*za makolo*) and improved maize (*za chizungu*). Several points were raised under this component. A table below (Table 3) provides a summary of what farmers ‘felt’ about local and improved maize.

**Table 3.** Farmers’ perceptions on local and improved maize during focus group discussions in Salima Agricultural Development Division, Salima District, Central Malawi from 2004/05-2018/19 period

| Why local maize? | Why not local maize? | Why improved maize? | Why not improved maize? |
| --- | --- | --- | --- |
| Good taste | Late maturing | High yielding | Easily attacked by storage and field pests |
| Retains more flour after milling | Low yield | Early maturing | High storage costs |
| Requires less flour for ‘nsima’ | Grows slowly | Drought tolerant | Less flour after milling |
| Pest resistance in storage | Visually small cobs | Some have recyclable seed | High flour demand for ‘nsima’ |
| Low storage costs |  |  | Tasteless |
| Cheap |  |  | No appetizing smell when cooked fresh |
| Direct seed source |  |  | Some not recyclable seed |
| Appetising smell when cooked fresh |  |  |  |

Participants highlighted that despite preferences, sometimes they just grew the maize they did not want. This applied to both either local or improved on one hand and varieties within improved maize. This component revealed wider views and needs of farmers on maize types as indicated by the statements below:

> ‘I sometimes grow improved maize because extension officers advocate for hybrids. They (extension officers) ‘admonish’ (*amatinena*) us when they find local maize in our fields. They say that the focus of the ministry of agriculture is on hybrid or any other improved maize types for higher yields’. [Local maize farmer)

> ‘I would not mind the type, whether local or improved provided the variety met what I wanted’. [Hybrid and local maize farmer]

Farmers also showed high regard for hybrid maize, especially on high yielding. They mentioned varieties like DK 8033, DK 8053, DK 777, MH 41, ZM 523 as exceptionally high yielding. They opined that hybrid maize could be better for dealing with incessant droughts experienced in the study area. They also indicated that SC 403 was popularly grown because it matured early, however, the variety was disliked because storage pests like weevils attacked it in the field before it would be ready for harvesting. They also indicated that the variety produced small cobs. The participants argued that for purposes of food security reasons, the prices of improved maize should be reduced so that all farmers afford them. One form of admiration for hybrid maize was demonstrated in the questions below:

> ‘Why is it not possible for researchers to maintain most of the good characteristics of local maize in hybrid varieties apart from high yielding and drought resistance?’ [Lead farmer]

### Special focus on poundability

A special focus on poundability was meant to understand some real perceptions, especially from women. This component brought interesting arguments. Participants indicated that improved maize especially hybrid maize had poor poundability and provided less flour. This brought in some quantitative measurement analysis among the participants. Some opined that for food purposes, a household with local maize would be food secure than with hybrid maize if they used the same annual consumption rate of 300 kg per capita. The following analogy was weighed in:

> ‘For those who grow maize for food, it is better to grow local maize. If you take three 50 kg bags of local maize to a mill and you would like to process ‘white flour’, you will come back with almost the same 3 bags as flour. But if you take the same number of bags of hybrid maize for the same purpose you will come back with only 1.5 bags (50% less) as flour and chaff/bran have a ratio of 1:1 for hybrids. We lose a lot. Unfortunately, some of us do not keep livestock to feed on the chaff. We just leave it with mill owners’. [Hybrid and OPV maize grower]

> ‘…….. I use more flour from hybrid maize to make ‘nsima’ compared to when I use flour from local maize’ [Local and hybrid maize farmer]

### Promotion priority between local and improved maize

This also formed a fiery debate among the farmers as they kept on shifting from local to improved maize but finally, a general understandable direction was drawn. Participants opined that all maize types should be promoted. They, however, indicated that with prevailing poverty levels among most rural communities, hybrid and OPV maize should not be sold at high prices. This was in light that many hybrid varieties (and OPV) were high yielding and therefore, suitable for hunger prevention. The participants acknowledged the biased promotion of improved maize over local maize. This was attributed to two big national government programs; the Agriculture Sector-Wide Approach-Support Project 1 (ASWAP-SP 1) from 2008/09-2013/14 which promoted hybrid and OPV maize variety performance through demonstrations and the Farm Input Subsidy Programme (FISP) from 2005/6 to present which subsidized farm inputs including hybrid and OPV maize seed. They indicated that many demonstrations were mounted where various hybrid and OPV maize varieties were compared for farmers to choose. The aim was for farmers to choose the best performing improved varieties in their area. The variety of performance demonstrations focused on maturity duration and yield. There were no local varieties in such demonstrations. On OPV and hybrid maize, farmers narrated that many FISP beneficiaries opted for OPV around the first three years because there was less top-up amount compared to hybrid varieties. But when they compared yields, they shifted to hybrid varieties. According to some farmers, this contributed to the declining focus on local maize. The participants opined that most farmers reduced land sizes under local maize. A statement in support of the diversity of maize was uttered:

> ‘It depends on what one wants if you want to meet quantity go for hybrid or OPV, but if you want quality go for local maize. It is good we have many choices.’ [Hybrid maize farmer]

> ‘When you grow improved maize, you are treated as s progressive farmer while local maize is associated with ‘limited knowledge’ in farming.’ [Hybrid and OPV maize farmer]

Direct Field Observations (DFO) confirmed that some farmers grew hybrid, OPV, local, and a combination of these maize types. Of the 10 fields that were visited, six were dominated by hybrid and OPV while three had both local and hybrid (with clear demarcation) and one had local only.

### Extent of local, hybrid, and OPV maize cultivation

From 2004/05-2018/19, the area under local maize and OPV decreased from 21 619 ha to 8 331 ha, and 12 913 ha to 11 309 ha respectively, while the area under hybrid maize increased from 9 811 ha to 19 379 ha (Figure 2A). This means that the area under local and OPV maize reduced by 61% and 12% respectively, while that of hybrid maize increased by 49%. Production and yield (Figure 2B and 2C respectively) for hybrid maize were very variable from 2014/15-2018/19 seasons.

**Figure 2.**
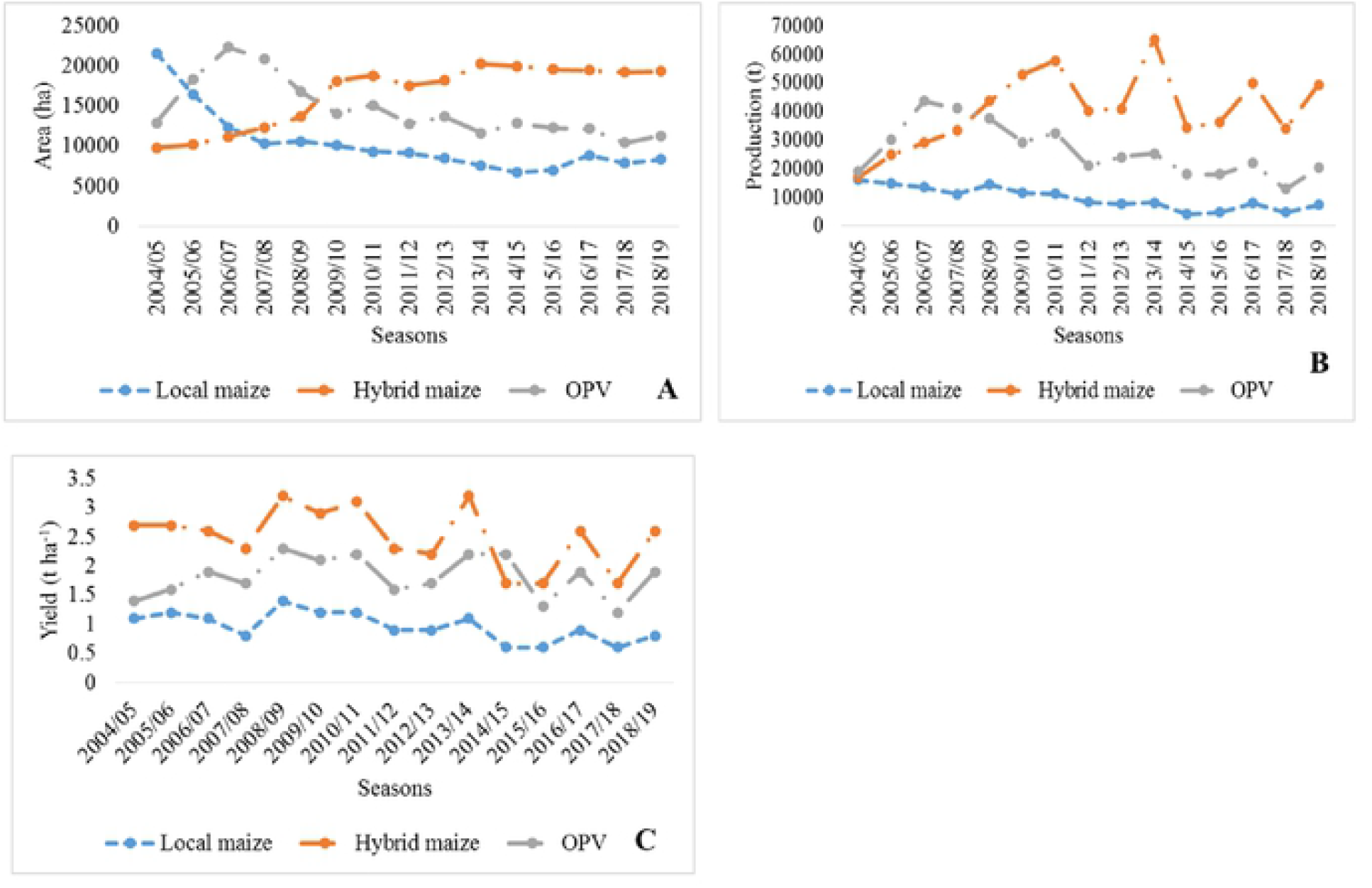
Trend of area cultivated (A), production (B) and yield (C) of local, hybrid, and OPV maize in Salima Agricultural Development Division, Salima District, Central Malawi from 2004/05-2018/19 Seasons

Over the study period, hybrid and OPV maize had been cultivated on a significantly larger area than local maize (Table 4). As expected the yield and production for the hybrid were higher than those of local and OPV maize. OPV maize yield was higher than that of local maize. However, out of the potential yield range of 7-10 t ha^−1^ for hybrid maize, the average yield was far too low at about 2.50 t ha^−1^. Taking 7 t ha^−1^ as the minimum yield for hybrid maize, it would be expected to realise production of about 115 696 tons on 16 525 ha during the period of the assessment but only 40 666.5 tons were realized. This represents 35% of the minimum potential production. For local maize, only 9 756.4 tons out of a potential of 30,000 tons were realized, representing 31%. Under OPV, only 26 352.5 tons were realised out of a potential of 101 664.9 tons on 14 520.7 ha representing 26%. Across the years, yields (Figure 2C) were lowest for hybrid maize in 2014/15 and 2015/16 seasons (1.7 t ha^−1^) when local maize had 0.68 and 0.62 t ha^−1^, and OPV had 2.2 and 1.3 t ha^−1^ respectively. Local maize had the lowest yield in 2017/18 (0.61 t ha^−1^) when hybrid and OPV maize had 1.8 and 1.2 t ha^−1^ respectively. All the maize types had yield peaks in the 2008/09 season and hybrid maize attained a similar yield in the 2013/14 season. The mean yield for all maize types was 1.71 t ha^−1^. Notable yield decrease was observed for local (51%) and hybrid (47%) maize from 2014/15-2015/16 and for all types in the 2017/18 season. In the 2017/18 seasons, hybrid, local, and OPV maize yields decreased by 45%, 55%, and 46% respectively.

**Table 4.** Comparison on average area cultivated production and yield of hybrid, local, and OPV maize in Salima Agricultural Development Division, Salima District, Central Malawi from 2004/05-2018/19 Seasons. Values with the same letters along a column are not significantly different. SD = Standard deviation, LCL = Lower confidence level, UCL = Upper confidence level

| Parameter | Mean area (ha) | SD | LCL | UCL | Minimum | Maximum |
| --- | --- | --- | --- | --- | --- | --- |
| Hybrid maize | 16 528a | 3 885.87 | 14 544.5 | 18 511.6 | 9 811 | 20 310 |
| Local maize | 10 304b | 3 960.42 | 8 340.80 | 12 307.9 | 6221 | 21 619 |
| OPV maize | 14 520.7a | 3 561.96 | 12 537.2 | 16 504.3 | 10 415 | 22 309 |
| t. statistic | 2.018 |  |  |  |  |  |
| P value | 0.000218*** |  |  |  |  |  |

| Mean production (tons) |  |  |  |  |  |  |
| --- | --- | --- | --- | --- | --- | --- |
| Hybrid maize | 40 666.5a | 12 869.72 | 35 774.47 | 45 558.59 | 17 180 | 65 371 |
| Local maize | 9 756.4c | 3 866.25 | 4 864.34 | 14 648.46 | 4 157 | 16 107 |
| OPV maize | 26 352.5b | 9 157.37 | 21 460.47 | 31 244.59 | 12 832 | 43 773 |
| t. statistic | 2.018 |  |  |  |  |  |
| P value | 0.0001*** |  |  |  |  |  |

| Mean yield (t ha<sup>-1</sup>) |  |  |  |  |  |  |
| --- | --- | --- | --- | --- | --- | --- |
| Hybrid maize | 2.50a | 0.5140 | 2.2982 | 2.7017 | 1.7 | 3.2 |
| Local maize | 0.97c | 0.2501 | 0.7583 | 1.1617 | 0.6 | 1.4 |
| OPV maize | 1.81b | 0.3502 | 1.6116 | 2.0150 | 1.2 | 2.3 |
| t. statistic | 2.018 |  |  |  |  |  |
| P value | 0.0001*** |  |  |  |  |  |

Figure 3 shows annual percentage changes in the area, production, and yield for all the three maize types. The percentage annual decrease in area under local and OPV maize was not significantly different from the percentage increase in area for hybrid maize (p = 0.556). For local maize, the net annual change in area and production was negative while that of yield was positive. The percentage annual changes in yield (p = 0.953) and production (p = 0.113) between local, hybrid, and OPV maize were not significantly different. Hybrid and OPV maize had positive net annual changes in all three parameters. During the 15 year period, rainfall variations significantly influenced the yield for hybrid (p = 0.03075; F = 4.719; *R*^2^ = 35%) and local maize (p = 0.00029; F = 17.3; *R*^2^ = 70%) and not OPV maize (p = 0.1484; F = 2.246; *R*^2^ = 15%) although, the coefficient of determination for OPV was small.

**Figure 3.**
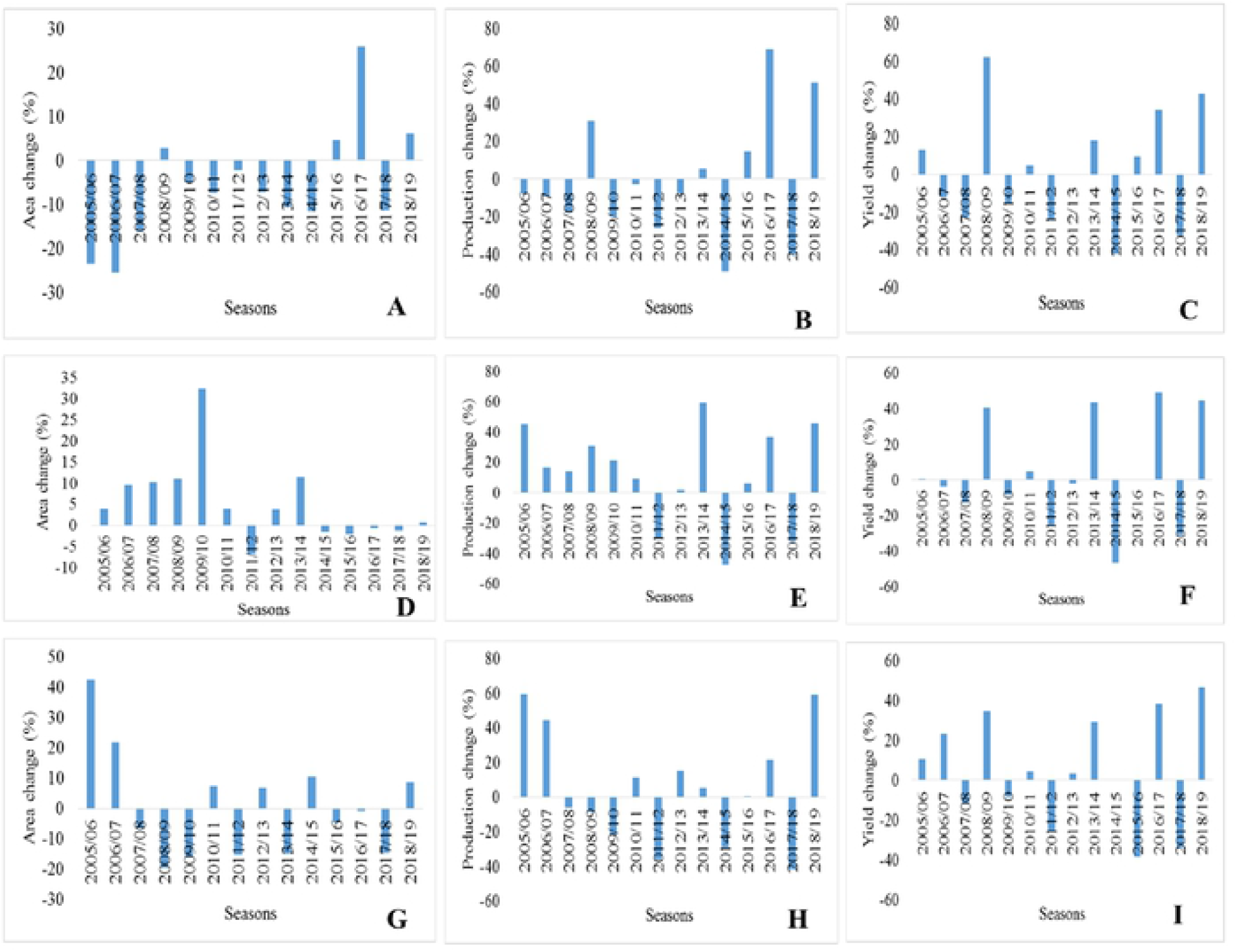
Annual percentage changes for area, production, and yield for local maize **(A-C)**, hybrid maize **(D-F)**, and OPV maize **(G-I)** in Salima Agricultural Development Division, Salima District, Central Malawi from 2004/05-2018/19 growing seasons.

An important observation was noted on the consistent and significant percentage increase (p = 0.0376) for the area under local maize from 2014/15 to 2018/19 and a decrease in the area for hybrid and OPV maize (Figure.4) in the same seasons. However, the percentage change in production and yield was not significantly different in these five years. The area under local maize increased by 403.9 ha annually while the area under hybrid and OPV decreased by 160.8 ha and 499.4 ha respectively. Production for local and hybrid maize increased annually while that of OPV decreased. Yield followed the same trend. Area for all maize categories decreased while production and yield for all maize combined slightly increased.

**Figure 4.**
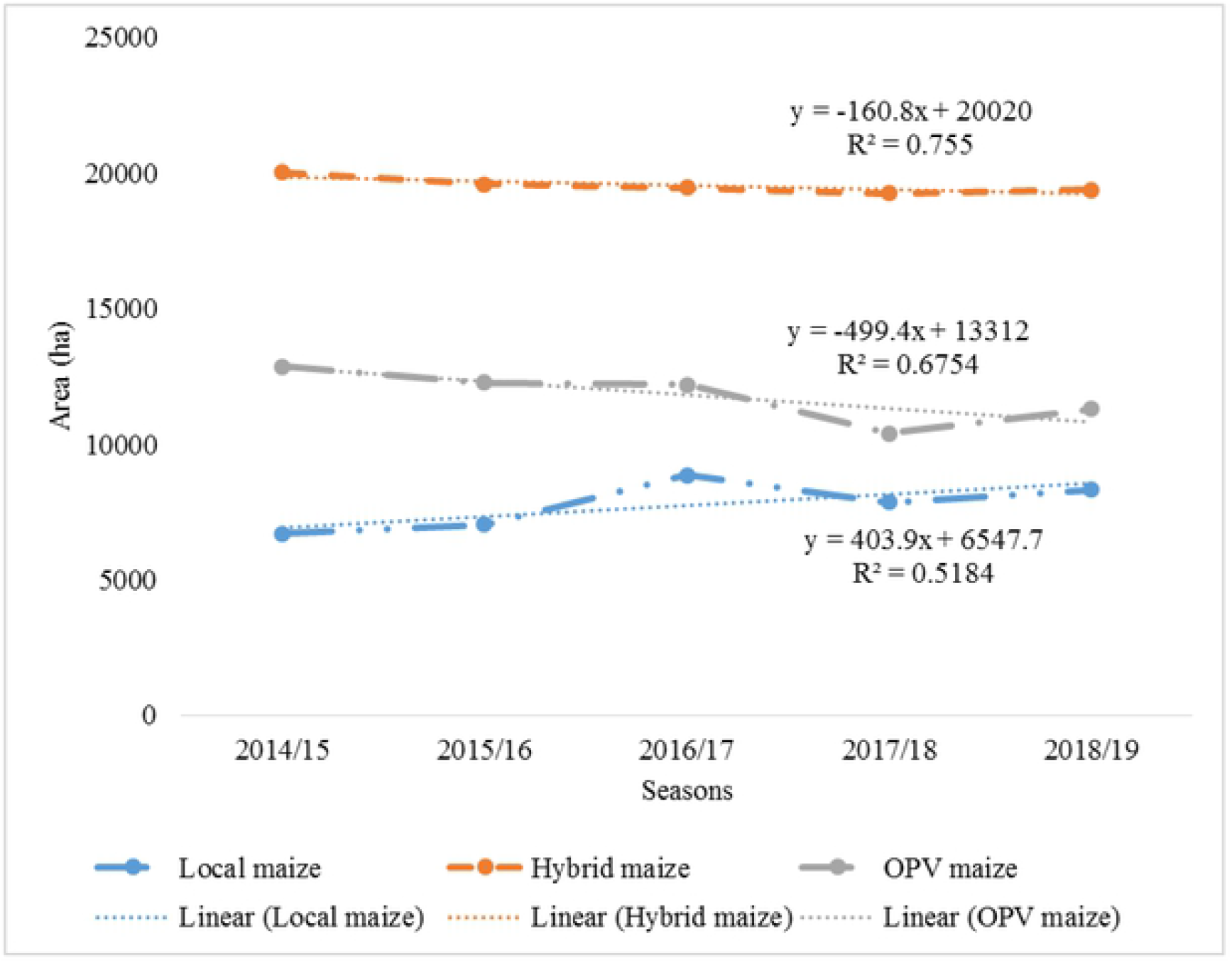
Annual change for the area under local, hybrid, and OPV maize in Salima Agricultural Development Division, Salima District, Central Malawi from 2014/15-2018/19 growing seasons.

### The outlook of all maize subsector in Salima (2019/20 to 2025/26)

Production of maize in Salima depends on several factors ranging from uncertain and uncontrollable factors such as rainfall and other natural calamities to factors such as the input costs, price of maize grain commodity, and economic growth. The study projected the future production of maize up to 2025/26 and the projections revealed that the district would experience a slight increase in thousands of tons of maize commodity during the period 2019/20 and 2025/26. The production of maize for Salima district is projected to be 44 172 metric tons by 2025/26 season, an increase of 1.6% from the base year of 2019/20 (Table 5). The production of maize will increase due to favorable climatic conditions and not necessarily by the increase in area and yield.

**Table 5.** Maize production trend as projected from 2019/20-2025/26 in Salima Agricultural Development Division, Salima District, Central Malawi.

| Season | The area under maize production (ha) | Maize yield (tons/ha) | Maize production (tons) | Area harvested growth rate (%) | Yield growth rate (%) | Production growth rate (%) |
| --- | --- | --- | --- | --- | --- | --- |
| 2019/20 | 89,629 | 2.06 | 43,467 | 4.10 | 13.92 | 10.23 |
| 2020/21 | 88,667 | 2.03 | 43,651 | -1.51 | -1.08 | 0.42 |
| 2021/22 | 87,539 | 2.00 | 43,752 | -1.52 | -1.29 | 0.23 |
| 2022/23 | 86,405 | 1.97 | 43,854 | -1.55 | -1.31 | 0.23 |
| 2023/24 | 85,270 | 1.94 | 43,958 | -1.57 | -1.33 | 0.24 |
| 2024/25 | 84,133 | 1.91 | 44,064 | -1.60 | -1.35 | 0.24 |
| 2025/26 | 82,994 | 1.88 | 44,172 | -1.62 | -1.37 | 0.25 |

The model suggests that if maize yield can be improved by approximately 30% (Figure 5) from the current productivity levels, maize production levels can increase to 110,430 metric tons from the baseline value of 82,994 metric tons by 2025/26 harvesting season as revealed in the graph below.

**Figure 5.**
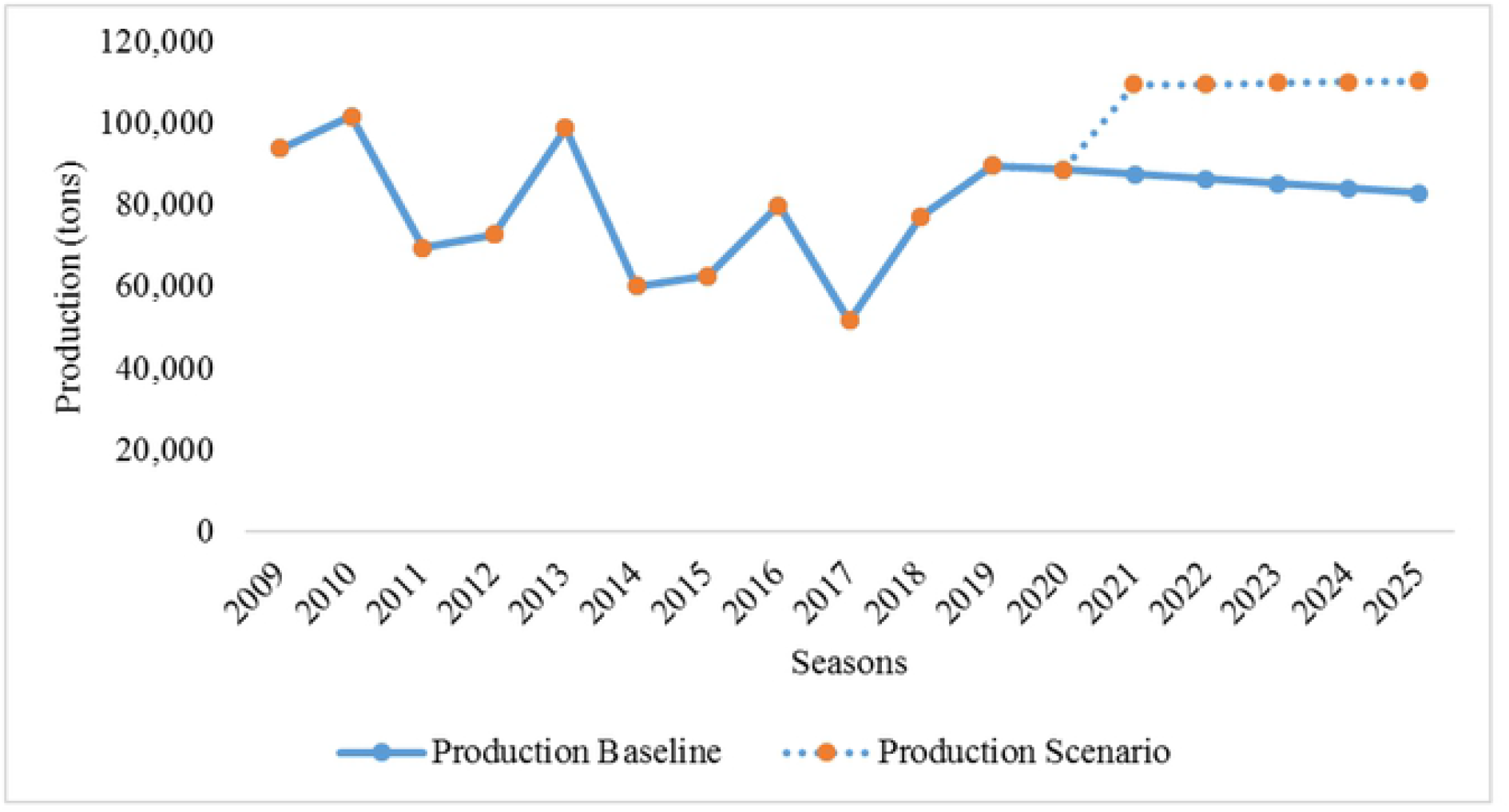
Projection of maize production under a 30% increase in yield from 2019/20-2025/26 in Salima Agricultural Development Division, Salima District, Central Malawi.

## Discussion

### Farmer’s perception of local, hybrid, and OPV maize

Here we discuss the maize varieties and factors farmers look for in a good maize type. Our results have shown that farmers had wider choices on over 15 hybrids, and six OPV maize varieties compared to two local varieties. The proliferation of hybrid maize varieties took an accelerated upward trend around the late 1990s where some hybrid varieties were distributed free of charge (Smale and Phiri 1998). Most of the improved varieties in our study had been reported in previous studies (Hoden and Fisher 2015). Farmers confirmed the dominance in the promotion of these varieties, especially from the late 2000s. The study revealed that only improved maize had formal markets dominated by Private Agrodealers. Where improved maize was under subsidy, farmers could either buy or collect free of charge from Private Agrodealers or ADMARC depending on the variety and prices of the seed. Our findings differ from earlier observations by Smale and Phiri (1998) that ADMARC was the main source of hybrid seed seconded by NGOs and relatives. This vindicates a shift in the proliferation of improved seed from a government institution to private traders. It was confirmed that no local maize could be found at any seed outlet. This could be due to policy directions which encouraged the use of improved maize (Government of Malawi 2006; Ministry of Agriculture, Irrigation and Water Development 2018). The sharing of local seed free of charge among farmers might mean that local maize is either undervalued or that its ‘low cost’ management (owned managed) makes it affordable.

The study has established ‘back to the drawing board’ insights for maize researchers, especially on attributes of good maize type and farmers’ perception of local and improved maize. The farmers’ top four preferences [1) High yielding, 2) adequate flour after milling, 3) cheap, 4) pest and disease resistance in field and storage], demonstrated that local maize was still valuable among local communities. Factors like adequate flour, cheap, and pest resistance in storage were associated with local maize more than improved maize. Farmers complained about hybrid maize for their susceptibility to storage pests, high storage costs, expensiveness, high flour demand to prepare ‘nsima’, and low flour after milling. This finding arguably confirmed that less progress might have been made in meeting farmer needs on hybrid maize varieties. Some past studies (Smale et al. 1991) observed that farmers in Kasungu (central Malawi), Blantyre (southern Malawi), and Mzuzu (northern Malawi) did not prefer hybrid because more flour would be needed to make ‘nsima’ than local maize while Pixley and Banziger (2001) found that smallholder farmers shunned hybrid maize due to high storage costs. Recently, Holden and Mangison (2013) chronicled ‘seed unavailability of hybrid maize’, ‘prefer local maize’, ‘no difference between local and hybrid maize’, ‘disease resistance’ as reasons for farmers’ preference for local maize. Farmers indicated that most improved maize varieties lacked ‘appetising smell when cooked fresh’. This came as a unique observation.

In this study, farmers appreciated the high yields realised from improved maize but still emphasized that some desired features of maize do not reflect in improved maize especially, hybrids. This could mean that the high adoption of improved maize and increase in area from 2005/06-2014/15, was accelerated because of the cheap subsidised inputs. Farmers might have been attracted by almost free hybrid and OPV maize seed. Based on the focus group discussion (FGD) and as observed by Dorward and Chirwa (2013) the reduction in FISP beneficiaries, especially from 2014/15-2018/19, might have led to a corresponding decrease in area under hybrid and OPV and an increase under local maize during the period (Figure 4 above). Non-FISP beneficiaries might have been failing to buy and meet the cost of hybrid and OPV maize production, hence opting for local maize. This resonated positively with the findings from FGD where farmers indicated ‘cheap’ as an important element for them to go for a variety. Salima district had 23 440 beneficiaries in 2005/06 and 18 000 in 2018/19 (Salima Agricultural Development Division 2019). Smale et al. (1991) and Ali et al. (2020), observed that farmers not only focus on the biological/physiological performance of a maize variety but also affordability. Farmers who grow maize for food usually go for local variety if they know that they would not meet the cost of production for hybrid maize (Machiria et al. 2010; Holden and Mangison 2013). In Malawi, farmers would prefer hybrids or OPV over local maize only if hybrid seed and fertilizers were highly subsidised (Denning et al. 2009). Therefore, it is important to focus on net valuation rather than mere yield (Smale et al. 1991). Yield advantage will only be deemed important if it translates into competitive socioeconomic benefits for farmers (Machiria et al. 2010). It would be speculated that the current dominance of private traders over hybrid maize seed could mean continued expensiveness of the seed for most Malawian farmers.

The FGD revealed that some farmers still did not know some differences between hybrid and OPV maize. Some could generally say that ‘*za chizungu*’ (improved maize) could be recycled while others said ‘*za chizungu*’ could not be recycled in terms of seed. This knowledge gap has the potential to twist farmers’ decision-making for a maize variety especially if they are interested in recycling the seed. The other potential source of confusion was when some included varieties like DK 8033, DK 8053, and MH 41 as examples of drought tolerant. These varieties are non-drought tolerant (Hoden and Fisher 2015). This demands adequate extension education among farmers to enhance decision making on improved maize varieties.

### The extent of local, hybrid, and OPV maize production

By 2004/05, land under local maize was significantly higher than land under hybrid and OPV maize until 2007/08 season. Production and yield followed a similar trend. This is similar to studies done in the early 1990s to -mid-2000s (Smale et al. 1991; Heisey and Smale 1995; Smale and Phiri. 1998; Chirwa 2006) where about 62% of the land was under local maize. In 2004/05, production for local and hybrid maize was not significantly different despite significant differences in yield. This could be attributed to large areas under local maize which offset the higher yield of hybrid maize. However, after 2007/08, both yield and production for hybrid remained significantly higher than those of local and OPV maize. The unique higher yields for OPV maize compared to hybrid and local maize in 2014/15 and 2015/16 seasons, suggested that OPV maize were more drought-tolerant compared to hybrid and local maize. This was confirmed by the regression analysis of yield against rainfall. The seasons experienced moderate to the severe drought that affected 19 944 ha out of the 38 908 ha of all maize in the study area (Check Appendix) when the district registered 39% yield loss (Salima Agricultural Development Division 2019). This loss might have been highly contributed by hybrid maize since its yield was even below that of OPV for the first time (2014/15) throughout the study duration. This shows that farmers should opt for drought-tolerant OPV maize should there be a likelihood of droughts in the area. The lowest yields and production in the 2017/18 season for all maize types could suggest that they were all seriously prone to Fall armyworm attacks (See Appendix). The season experienced an outbreak of Fall armyworm that affected 19 839 ha out of the 37 511 ha of all maize. However, within the recent five years (2014/15-2018/19), the area under local maize increased by 26% with a corresponding production of 76%. The area under hybrid and OPV maize decreased by 3% and 12 % with a production increase of 44% and 13% respectively. This finding should arouse curiosity in the promotion or management of hybrid and OPV maize in Salima.

Although the yield for local maize (0.97 t ha^−1^) was lower than hybrid maize (2.50 t ha^−1^) and OPV (1.81 t ha^−1^) in this study, it was comparable to other studies of similar nature. Yield range of 1.3-2.0 t ha^−1^ for hybrid and 0.91-1.1 t ha^−1^ for local maize were reported in southern and central Malawi (Tchale and Sauer 2007; Holden and Fisher 2015). The yields in our study are a complete departure from yields from on-station research studies where up to 14.9 t ha^−1^ for hybrid and 11.0 t ha^−1^ for local maize were reported from Chitedze Research Station (Mubanga et al. 2018). Other studies have shown that under farmer conditions, local varieties performed better than hybrid varieties in terms of disease/pest resistance (Ebenebe et al. 2000; Mubanga et al. 2018), and non-use of fertilizer (Heisey and Smale 1995; Macharia 2010; Holden and Mangison 2013). Coincidentally, these were the factors that attracted farmers to opt for local maize as highlighted in the focus group discussions. Under farmer conditions, all necessary management measures should be addressed if all the maize types were to provide meaningful yields.

### Projected all maize production in Salima

Our study asserts that the use of hybrid and OPV maize alone based on high yielding may not be a panacea for food security under farmer conditions in Salima Agricultural Development Division. With the projected 2.6% annual increase in population (Government of Malawi 2018b), there is a need to emphasize high production per unit area of land which was not promising in the findings of this study. The study, therefore, suggests the need for production strategies that will enable the district to cope up with the population driven-demand for maize commodity in the next seven years. During the past years, increases in maize production in Salima were basically due to the expansion in area under cultivation. However, the maize yield level has historically been less than the potential levels, ranging between 1.2 to 2.5 tons ha^−1^. This, therefore, calls for government, the district agriculture office and other stakeholders to review the past interventions and policies in the maize subsector. The farmers ought to adopt management practices that could enable the district to increase not only the area under maize cultivation but also its productivity levels. High technical efficiency for maize production is maximized beyond the use of improved variety or chemical fertilizers but with the inclusion of integrated soil fertility and soil and water management interventions (Tchale and Sauer 2007; Ali et al. 2020). The study also found a positive response of area under cultivation to own and substitute crop prices. This implies that the producers in Salima can find it necessary to adjust the area allocated to maize production given changes in the prices of maize and rice as a second crop. The pricing strategies could bring about an effective and profitable adjustment in terms of sizes allocated to maize production or its alternative, i.e. rice

### Conclusions

The study aimed at understanding the local, hybrid, and OPV maize production under smallholder farmers in Salima Agricultural Development Division (SLADD), Central Malawi. Of interest, we focussed on establishing maize varieties (local, hybrid, and OPV) used, factors that farmers look for in preferred maize type or varieties, and revealed the extent of local, hybrid, and OPV maize cultivation in terms of area, yield and production in the long term from 2004/05-2018/19 and projected all maize production from 2019/20-2025/26. We adopted a Constructivism Epistemology (CE) approach through focus group discussion (FGD) to understand the perceptions of farmers on local, hybrid, and OPV maize. We also collected and analyzed maize production data for 15 years to understand the extent of maize production in SLADD.

Sixteen hybrids (SC 627 (*Mkango*), SC 403 (*Kanyani*), SC 719 (Njovu), SC 407, DK 9089, DK 8053, DK 8033, DK 777, MH 18, MH 19, MH 26, MH 31, MH 41, PAN 4M-19, PAN 53, Pannar 413), six OPV (ZM 421, ZM 521, ZM 523, ZM 621, ZM 623, Chitedze 4) and two local maize varieties (*Kaduya* and *Bantam*) were identified as commonly grown in the study area. Although sources of improved maize were numerous, farmers expressed a lack of such markets for local maize. Farmers used their stock for local maize. Farmers listed nine attributes of a good maize type or variety in the order as follows: 1) High yielding, 2) adequate flour after milling, 3) cheap, 4) pest and disease resistance in field and storage, 5) good taste, 6) good poundability, 7) early maturing, 8) demanding less or no fertilizer 9) readily available. Farmers preferred hybrid and OPV maize for high yielding and drought resistance and recommended them especially if the aim were to achieve the quantity of maize. Despite low yields some farmers still grew local maize because of good taste, cheap, adequate flour after milling, appetizing smell when cooked fresh, pest and disease resistance in field and storage, and good poundability which lacked in most hybrid and OPV maize varieties.

On the extent of maize production in SLADD, the area under local maize decreased by 61%, OPV by 12 % while the area under hybrid maize increased by 49% during the period of the study. Sharp decrease for the area under local maize and increase under hybrid maize were pronounced from 2008/9-2013/14 due to the promotion of hybrid maize through Agricultural Sector-Wide Approach-Support Project 1 (2008/09-2012/13) and the Farm Input Subsidy Programme (FISP) (2005/6 to present) as narrated during FGD. The increase in area for OPV from 2005/06-2006/07 was attributed to FISP. During the period under study yield for local, hybrid, and OPV were 0.97 t ha^−1^, 2.50 t ha^−1^, and 1.81 ha^−1^ with a production of 9 756.4 tons, 40 666 tons, and 26 352.5 tons respectively. The study established that under farmer conditions, only 26-35% of maize production potential could be achieved. When the findings from FGD (weaknesses and strengths of local and hybrid maize) and field production data were related, it would appear that some farmers were not fully convinced with the performance of hybrid maize varieties despite high yielding. They indicated that they grew hybrid to meet household food demand at the expense of other important qualities (taste, adequate flour, the good smell when cooked fresh). One of our important findings showed that there could be bottlenecks associated with the management and promotion of hybrid and OPV maize from 2014/15 to 2018/19 seasons causing the reduction in area under hybrid and OPV maize and increasing area under local maize. This may have serious food self-sufficiency impacts. We can argue that local maize will always be a ‘fall back on’ alternative should the management of hybrid and OPV maize fail to meet farmers’needs or remain expensive. Production of all maize was projected at 44 172 tons by 2025/26, representing a 1.6% increase from the base year 2019/20. The production of maize will increase due to favorable climatic conditions and not necessarily by the increase in area and yield. If the yield were improved by 30% within the projection period, the production would reach 110 430 tons representing 67% of the maize food requirement in the study area.

The findings of this study bring together scientists, policymakers, and extension experts on the need to rebuild strategies for meeting farmer needs and achieving food self-sufficiency. There is a need to ensure that some qualities associated with local maize i.e. good taste, the appetizing smell when cooked fresh, poundability, pest resistance in storage be imbedded in hybrid and OPV maize. The attainment of high yield in hybrid and OPV maize is still a far-fetched dream under conditions faced by smallholder farmers in Malawi. There is a need to incorporate other production means such as organic manure, fertilizer trees, soil, and water conservation technologies if we are to achieve meaningful yields with smallholder farmers. There is also a need to explore ways of supporting preferences for some farmers such as the establishment of stable markets for local maize as demanded by some farmers as it has been shown that hybrid and OPV maize were not a solution for meeting farmer needs at least in the short and medium terms. Local seed banks would be a progressive approach for meeting seed access for communities for local maize. Policymakers should also consider reducing the prices of hybrid and OPV maize if we were to enhance affordability by many rural farmers as some programs e.g. FISP did not target all farmers. Otherwise, the upward curving of the area under local maize in the recent five seasons and the projected decrease in production for all maize from 2019/20-2025/26 should be treated as a harbinger of an impending production crisis in the study area.

## Acknowledgements

The authors appreciate all the Agricultural Extension Development Officers (AEDOs) Agricultural Extension Development Coordinators AEDCs) for data collection throughout the period of study. All farmers are appreciated for their cooperation during data collection in their maize gardens. We thank Mr Gorge Bota of Salima District Agriculture Office for assisting in data reception and consolidation. The management of Agricultural Development and Marketing Corporation (ADMARC) for Salima District is appreciated for sharing information on maize and rice stocks and prices which were part of the data used in projecting maize production.

## Availability of data and materials

Data is available upon request from authors

## Authors’ contributions

HN conceived the study idea, design, data collection, data analysis and drafted the manuscript. He also coordinated the process of data collection from the district office. WM conducted data collection, data analysis and refined the manuscript. EMN refined the manuscript.

## Appendix

Major disasters noted with effects on the area under maize production during the period of the study in Salima Agricultural Development Division, Salima District, Central Malawi from 2004/05-2018/19 Seasons (the area and household affected are a combination of local, Open-pollinated varieties and hybrid maize)

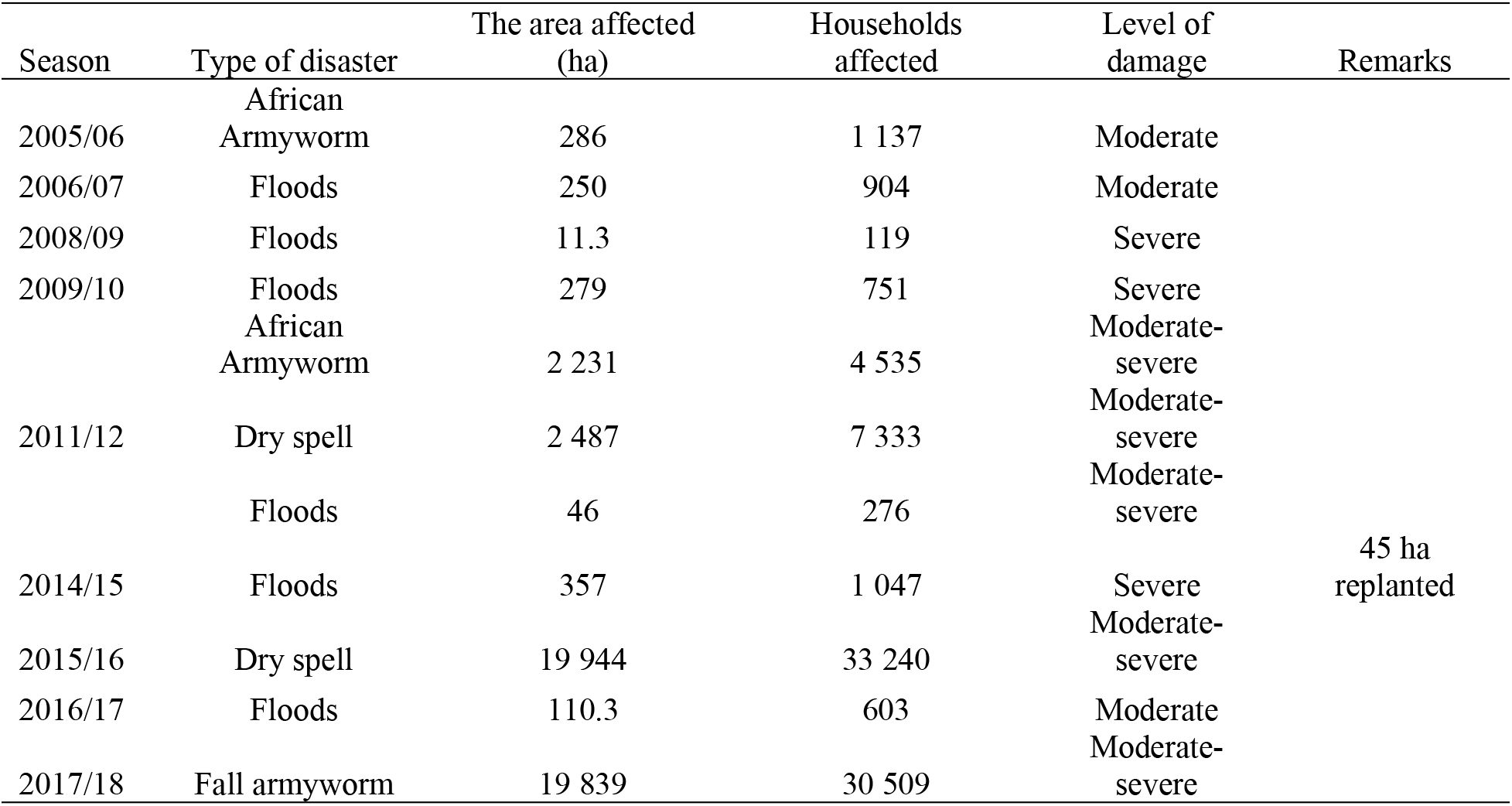

